# Chemosensory Adaptations in *Caenorhabditis* Males during the Establishment of Androdioecy

**DOI:** 10.1101/2025.05.24.655903

**Authors:** Harini Kannan, King L Chow

**Affiliations:** Division of Life Science, The Hong Kong University of Science and Technology, Clear Water Bay, Kowloon, Hong Kong, CHINA

## Abstract

*Caenorhabditis elegans* has evolved from its dioecious ancestors to adopt an androdioecious reproductive strategy. In this process, ancestral female *C. elegans* acquired genetic modifications that enabled self-sperm generation, self-sperm activation, and a reduced reliance on sexual reproduction. However, how males have adapted during this transition from dioecy to androdioecy is less explored. Using respective *Caenorhabditis* species, we demonstrated that androdioecious hermaphrodites exhibit a reduction in sex pheromone potency, while androdioecious males show notably heightened olfactory habituation and diminished mate exploration capabilities. The behavior of androdioecious males can be reverted to resemble that of dioecious males by replacing the SRD-1 receptor with its dioecious orthologs. This intrinsic characteristic is contingent upon the cytoplasmic domain of the receptor. We propose a theoretical framework where *C. elegans* males have accumulated genetic variations in their pheromone receptor, leading to altered chemosensory perception of the opposite sex, which confer a selective advantage that favors the establishment of hermaphroditism. Our study provides insights into an overlooked male trait that was shaped by changes in chemosensory signaling. The findings underscore the capacity of chemosensory variations to influence how organisms perceive critical ecological factors and eventually facilitate the emergence and stabilization of hermaphroditism.

## 1. Introduction

In the *Caenorhabditis* clade, evolutionary shifts have led to diverse mating strategies, with dioecy as the dominating mode.^1,2^ Mutations allowed ancestral females to produce self-sperm, enabling self-reproduction and reducing male reliance.^3–5^ Some lineages evolved into self-fertilizing hermaphrodites,^6^ exhibiting weaker sex pheromone potency^7^ and altered behaviors to avoid males^8,9^. Younger hermaphrodites resist male advances, while older and sperm-depleted ones show higher female-like receptivity.^10^ Hermaphroditism was thought to have evolved in low-density environments as an adaptive response to ecological pressure^11^. Social interactions also shape reproductive strategies, such as cooperative breeding in birds fostering monogamy or polyandry,^12,13^ underscoring the influence of social factors in mating system evolution.^12^ Historically, hermaphroditism research focused on a two-stage model of female-to-hermaphrodite transition, emphasizing self-sperm production and activation.^14,15^ However, male traits’ roles may have been overlooked. Our investigation elucidates male phenotypic characteristics associated with sex pheromones, including enhanced olfactory habituation and diminished mate exploratory behavior, thereby differentiating androdioecious males from their dioecious equivalents. The behavior of androdioecious males can be reverted to resemble that of dioecious males by replacing the SRD-1 receptor with its dioecious ortholog. This intrinsic characteristic is contingent upon the cytoplasmic domain of this receptor, implying that modifications in chemosensory signaling may have conferred a selective advantage to the establishment of androdioecy in *Caenorhabditis* species. Our study underscores the capacity of chemosensory variations to influence how organisms perceive critical ecological factors and eventually facilitate the emergence and stabilization of hermaphroditism.

## 2. Materials & Methods

### Worm strains

N2 CB4088:*him*□*5(e1490)V* was used for wildLJtype *C. elegans* for high incidence of males; EM464 (*C. remanei*), AF16 (*C. briggsae*); JU1428 (*C. tropicalis*); ZZY0401 (*C. sinica*); JU1904 (*C. wallecei*); CB4018:*fog*□*2(q71)* (*C. elegans* hermaphrodites lacks self-sperm); For experiments expressing SRD-1 orthologs in *C. elegans srd-1* mutant males we used KC301:*srd*□*1(eh1)II, him*□*5(e1490)V;* KC1445:*srd*□*1(eh1)II, him*□*5(e1490)V, wxEX[Psrd*□*1::srd*□*1:gfp unc*□*54 3*′*UTR + Pcrm1b::rfp unc*□*54 3*′*UTR];* KC1447:*srd*□*1(eh1)II, him*□*5(e1490)V, wxEX[Psrd*□*1::Cre*□*srd*□*1::gfp unc*□*54 3*′*UTR + Pcrm1b::rfp unc*□*54 3*′*UTR];* KC1449:*srd*□*1(eh1)II, him*□*5(e1490)V, wxEX[Psrd*□*1::Ce*□*srd*□*1*⍰*CT::Cre*□*srd*□*1 CT::gfp unc*□*54 3*′*UTR + Pcrm1b::rfp unc*□*54 3*′*UTR]*. **Sex Pheromone Extraction**

Synchronized L4 *C. remanei* and *fog-2* mutant *C. elegans* were separated, and after 24 hours, five virgin adult *C. remanei* (EM464) or 100 virgin *fog-2 C. elegans* females were incubated in 100 μl of M9 buffer at 25°C for 6 hours. The supernatant was then assessed on wild-type *C. elegans* or *C. remanei* males for quality control in the chemoattraction assay.

### Chemoattraction Assay

A microscopic slide was coated with 2 ml of chemotaxis agar (1.5% agar, 25 mM NaCl, 1.5 mM Tris-base, 3.5 mM Tris-Cl) and placed in a 5.5-cm Petri dish. Two 2 μl droplets of 1 M sodium azide were applied 3 cm apart and evaporated. Then, 2 μl of extract supernatant and control buffer were added to these spots. Twenty synchronized one-day-old adult males were placed at the slide’s center, equidistant (1.5 cm) from test and control sites. After 30 minutes, immobilized worms were counted based on their positions: C (control area), E (experimental area), or N (other regions). The Chemotaxis Index (CI) was calculated as CI = (E - C) / (E + C + N), with values near 1.0 indicating strong attraction and near zero showing no attraction. Each test included ≥20 assays with 20 males each, totaling 400 animals. Results are shown as CI ± SD, with significance assessed via one-way ANOVA with Bonferroni correction (***P<0.001).

### Mate Exploration Assay

Mating exploration was studied by placing a fixed number of *Caenorhabditis* males in a culture dish, allowing 10 minutes for dispersal, then introducing the same number of *C. remanei* virgin females to the dish. Observations lasted 30 minutes under a microscope, with vulval prodding indicating successful exploration. The earliest vulval prodding time on each plate was recorded to create a time vs. density graph. Population density was altered by adjusting dish sizes (3.5 cm, 5 cm, 10 cm for high, medium, and) and animal numbers. Each test included 15 assays, and significance was assessed using one-way ANOVA (***P<0.001).

## 3. Results and Discussion

The transition to androdioecy from dioecious progenitors is not exclusive to *Caenorhabditis* species;^16^ similar such transitions have occurred in *Datisca glomerata*^17,18^ and *Eulimnadia texana*.^19^ Androdioecy is advantageous in ecological contexts with high rates of extinction and recolonization.^20^ *C. elegans*, recognized for its fluctuating population dynamics experiences a persistent “boom and bust” cycle^21^ where androdioecy adoption may be an adaptive strategy to mitigate extinction risk. *C. elegans* males exhibit lower sex pheromone sensitivity than *C. remanei* males,^10,22^ suggesting that gonochoristic *Caenorhabditis* males could be more efficient in mate pursuit.

### Androdiecious males exhibit deficient mate exploration and strong sex pheromone habituation

Using *C. elegans* and *C. remanei* males as a proxy for the respective mating systems in this study, we assessed the mate-seeking abilities of androdioecious and gonochoristic males. Using light microscopy, we measured the time males took to locate virgin females and documented mating behaviors, such as vulval prodding,^23^ within 30 minutes. *C. remanei* females were chosen for their ability to attract males across *Caenorhabditis* species.^10^ Mating behaviors under different population densities at a 1:1 male-to-female ratio were evaluated. In a 10 cm culture dish with 20 androdioecious males and 20 virgin *C. remanei* females, androdioecious males took over 25 times longer to initiate copulation than dioecious males.

Fifteen or ten or five androdioecious males against equal number of *C. remanei* females results in failure of locating the females within the 30 min period, while dioecious males took 1 ± 0.8, 4 ± 1.35 and 4 ± 2.13 minutes respectively to do so (Fig 1B). When only one pair of animals were tested, neither group mated within 30 min. When the same set of experiments were repeated in medium and higher density conditions in 5cm and 3.5cm dishes, improved performance of androdioecious males was observed, although dioecious males consistently outperformed their androdioecious counterparts and initiated active mating within a shorter period irrespective of their population density (Fig 1B, grey and black bars). Inadequate mate exploration was identified in other androdioecious *Caenorhabditis* species as well (Sup Fig 2A), whereas dioecious *Caenorhabditis* species exhibited consistent and proficient mate searching behavior (Sup Fig 2B). Although androdioecious males can locally search and copulate with females or hermaphrodites in proximity (Kannan and Chow, unpublished), their prolonged mate location time may indicate a deceptive perception of prospective partners nearby, as if they were in a habitat with low-population density.

**Figure 1:**
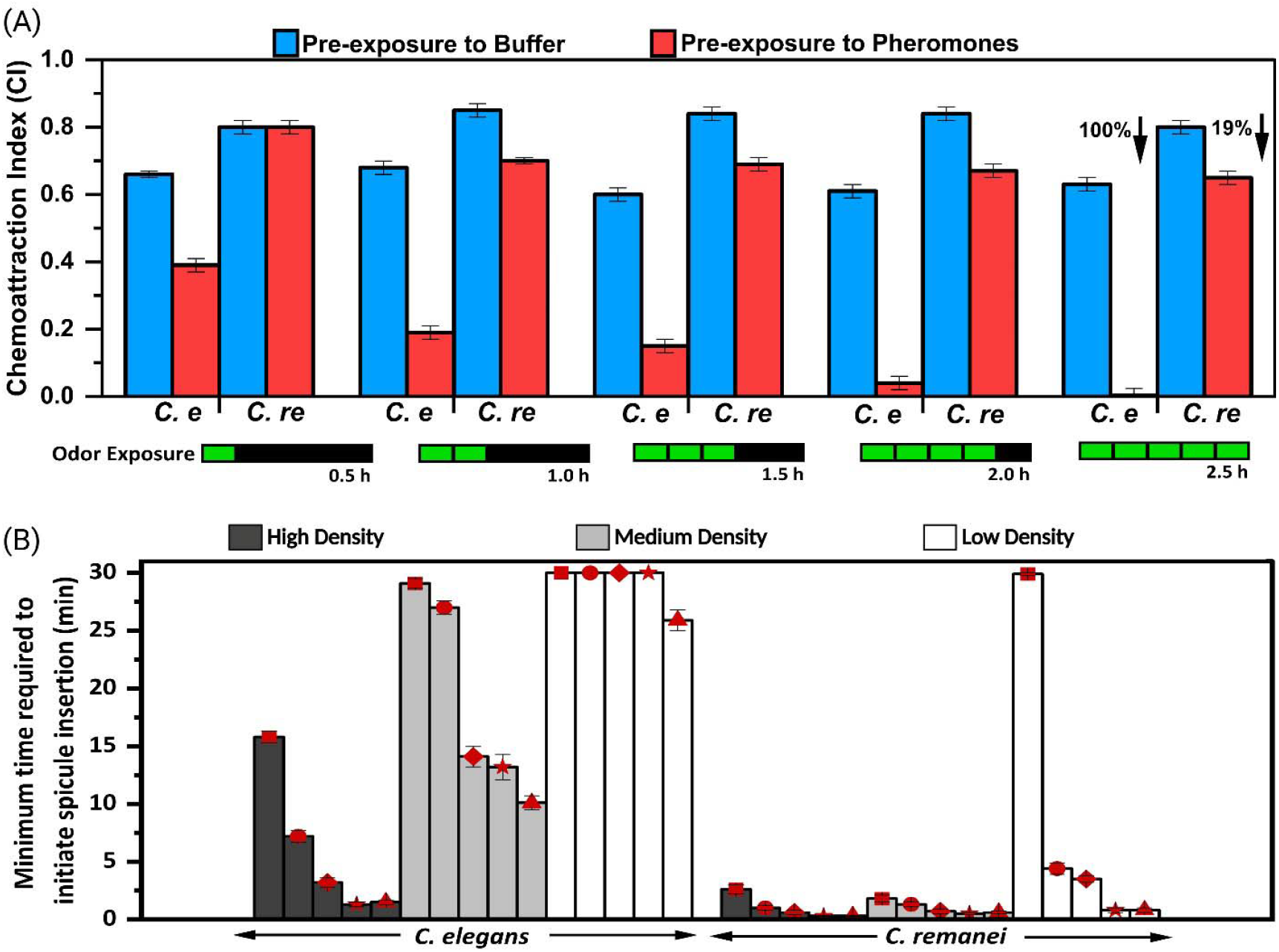
**(A)** Olfactory habituation was evaluated by measuring the response of *C. elegans* and *C. remanei* males to *C. remanei* sex pheromones following pre-exposure to either pheromones or M9 buffer (*n* = 400); (B) The minimum time required for *C. elegans* and *C. remanei* males to initiate vulval prodding on *C. remanei* females was recorded under varying density conditions (symbols: square = 1 pair, circle = 5 pairs, diamond = 10 pairs, star = 15 pairs, triangle = 20 pairs; *n* = 15). High, medium, and low-density tests refer to experiments conducted in 3.5cm, 5cm and 10cm culture dishes.

Signal transduction components essential for *C. elegans* to detect sex pheromone cues overlap with sensory pathways that perceive nutritional signals;^24–26^ prolonged exposure to these signals induce olfactory habituation, thereby diminishing the animals’ sensitivity to nutritional stimuli.^27^ To determine if sex pheromones trigger similar habituation phenomena and whether patterns differ between species of different mating types, we exposed androdioecious (*C. elegans, C. briggsae, C. tropicalis*) and gonochoristic males (*C. sinica, C. remanei, C. wallecei*) to *C. remanei* sex pheromones for varying durations. We then measured their chemoattractive indices (C.I.) in response to the pre-exposed odor (sex pheromones) in contrast to the control group exposed to M9 buffer. Androdioecious males showed the strongest habituation with their responses drastically compromised after 2.5 hours of sex pheromone pre-exposure: *C. elegans* (CI: 0.003 ± 0.07) (Fig 1A), *C. briggsae* (CI: 0.2 ± 0.09), *C. tropicalis* (CI: 0.005 ± 0.06) (Sup Fig 1A). In contrast, gonochoristic males remained highly responsive: *C. remanei* (CI: 0.65 ± 0.09) (Fig 1A), *C. sinica* (CI: 0.60 ± 0.1), *C. wallecei* (CI: 0.602 ± 0.1) (Sup Fig 1B). Impaired mate-searching ability and robust sex pheromone habituation are distinctly observed in contemporary androdioecious males likely having offered them selection advantages while also differentiating them from their dioecious equivalents.

### Hermaphrodites continue to produce sex attractant but at a lower potency

Sex pheromones from dioecious females are more potent than the attractants from hermaphrodites. Pheromone extracts from 5 one-day-old virgin *C. remanei* females attracted conspecific males and males from various other species (Sup Fig 4D), while extracts from 5 one-day-old *C. elegans* N2 hermaphrodites failed to elicit male response (Sup Fig 4C). Lower concentrations or chemical differences may explain the weaker allure of hermaphrodites, challenging the idea that they lack pheromones entirely.^7,10^ To verify this notion, we extracted pheromones from different numbers of *C. elegans* N2 hermaphrodites. Pheromones from 100 one-day-old *C. elegans* N2 hermaphrodites were about half as attractive (CI: 0.31 ± 0.1) as those from 5 virgin *C. remanei* females (CI: 0.72 ± 0.08) (Sup Fig 4C). Since hermaphrodites differ from females primarily by their ability to produce self-sperm, we further tested whether presence of sperms could impact the attractiveness. Using *C. elegans fog-2* mutants, which produce pseudo-females lacking self-sperm,^28,29^ we found pheromone extracts from 5 to 40 pseudo-females were unattractive to *C. elegans* males (CI: <0.20 ± 0.07) while extracts from 100 pseudo-females (CI: 0.72 ± 0.07) matched the potency of those from 5 *C. remanei* females (Sup Fig 4C). If concentration differences explain this disparity, female-derived pheromones should exceed the threshold for robust attraction. Titration of *C. remanei* pheromones showed attractiveness (CI: 0.49 ± 0.08) persisted even after a 1000-fold dilution (Sup Fig 4E). GC-MS analysis of pheromones from *C. remanei* females and *C. elegans* hermaphrodites revealed similar chemical profile with a key molecule contributing significantly to potency in both blends (Chan and Chow, unpublished). The weaker sex appeal, compared to *C. remanei* females, is primarily due to reduced pheromone potency. These findings contradict the earlier reports that *C. elegans* hermaphrodites have lost the sex pheromone production ability.^7,10^

### The chemosensory traits observed in androdioecious *Caenorhabditis* males converge on SRD-1

Apart from conservation in the sex pheromone blend, con-specific attraction elicited by female and hermaphrodite sex pheromones among *Caenorhabditis* males (Sup Fig 4D) may also be ascribed to the conservation of the sex pheromone receptor expressed in males. The amino acid compositions of receptor orthologs from *C. elegans* and *C. remanei* SRD-1 exhibited high conservation within their N-terminal region with notable polymorphisms primarily located in their cytoplasmic domain (Sup Fig 4A). Similar observation was noted in SRD-1 orthologs across various *Caenorhabditis* species, where N-terminal regions are highly conserved and distinct polymorphic variations were identified in the cytoplasmic domain (Sup Fig 3, Sup Fig 4A). Notwithstanding the polymorphism observed within the cytoplasmic domain, the SRD-1 ortholog from *C. remanei* demonstrated an effective functional integration with the sex pheromone sensory circuitry of male *C. elegans. C. elegans srd-1* mutant males, when expressing *C. remanei srd-1* transgenes regulated by the 3 kb long native *srd-1* promoter, resulted in a restoration of sex pheromone perception that was marginally more pronounced than that observed in *C. elegans srd-1* mutant males expressing the native receptor (Sup Fig 4B). Assessments conducted on 3-4 independent transgenic lines suggest that the observed phenotypic traits were not confounded by variations in transgene copy number. Conservation in the sex pheromone communication mechanism among androdioecious and dioecious *Caenorhabditis* species, at the pheromone, its receptors, and the functional integration in androdioecious males, prompted us to test whether substitution of the sex pheromone receptor would alter male sexual behavior.

We replaced the *srd-1* gene of *C. elegans* males with the dioecious ortholog from *C. remanei* and evaluated the habituation trajectory with *C. remanei* sex pheromones and mate exploration pattern of the male subjects against 1-day-old virgin *C. remanei* females. After 2.5 hours of pheromone exposure, *C. elegans srd-1* mutant males expressing the native SRD-1 completely lost the sensory response loss (CI: -0.01 ± 0.07), mirroring wild-type *C. elegans* (Fig 2A). In contrast, those expressing *C. remanei* SRD-1 retained ∼60% responsiveness (CI: 0.35 ± 0.09) (Fig 2B). Due to significant polymorphisms between *C. elegans* and *C. remanei* SRD-1 residing in their cytoplasmic domain, we created a chimeric receptor by substituting the *C. elegans* SRD-1 cytoplasmic domain with that of *C. remanei* SRD-1. *C. elegans srd-1* mutant males expressing the chimeric receptor retained their sensory response (CI: 0.44 ± 0.08) after 2.5 hours of pheromone exposure resembling those expressing *C. remanei* SRD-1 orthologs (Fig 2C).

**Figure 2:**
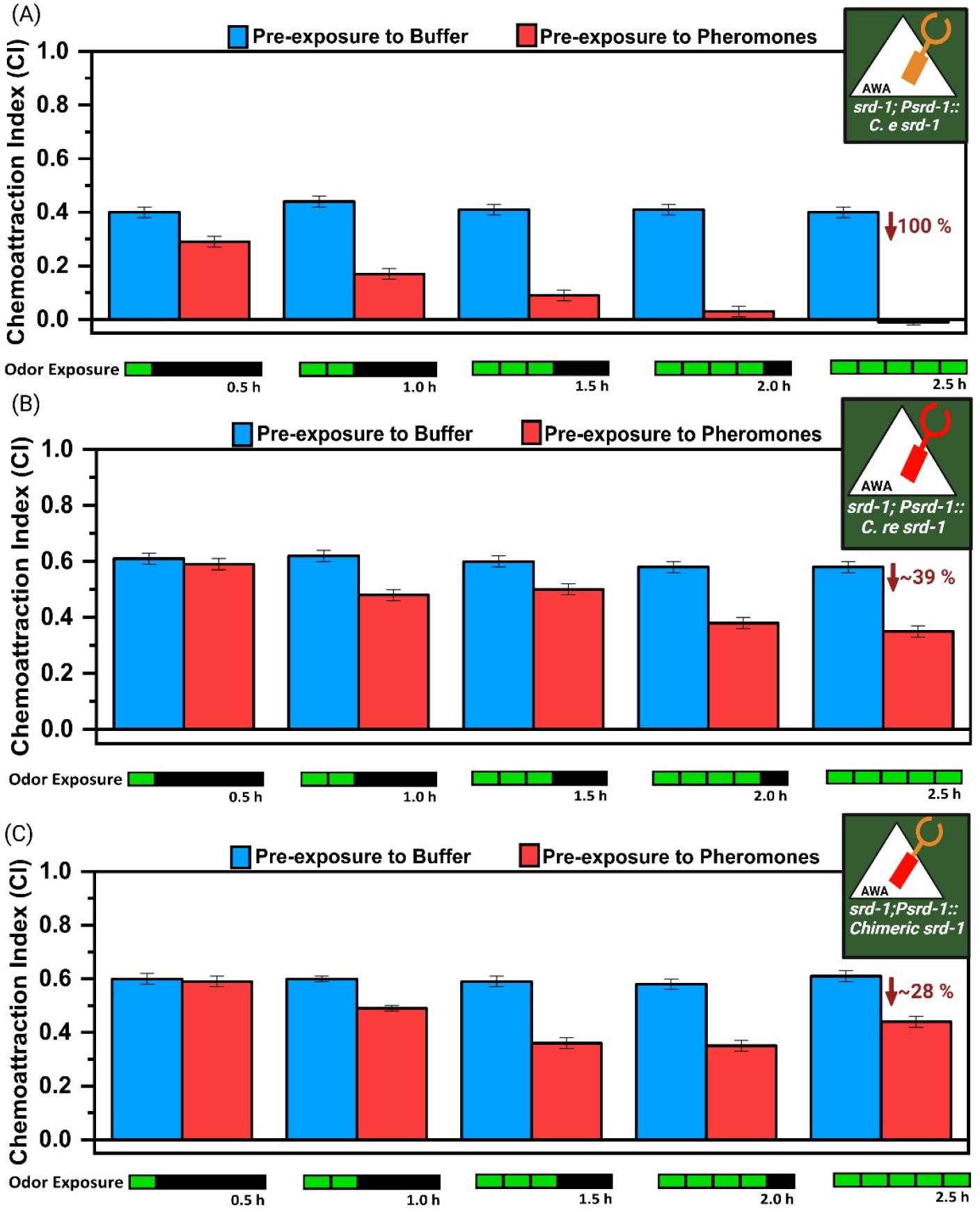
Olfactory habituation was assessed by measuring responses of *C. elegans srd-1* mutant males expressing (A) *C. elegans* SRD-1, (B) *C. remanei* SRD-1, and (C) Chimeric SRD-1 to *C. remanei* sex pheromones after pre-exposure to pheromones or M9 buffer (*n* = 400 males).

We further investigated the influence of SRD-1 on the animals’ ability to locate and participate in copulation. In a 10 cm culture dish containing 20 pairs of males and females, *C. elegans* transgenic males expressing *C. elegans* SRD-1 took an average of ∼26 ± 3.8 minutes to initiate mating. In contrast, those expressing *C. remanei* SRD-1 and chimeric SRD-1 averaged ∼3 ± 1.2 and ∼5 ± 1.7 minutes, respectively (Fig 3A, B & C). When the number of animals were reduced to 15 or fewer pairs, transgenic males expressing *C. elegans* SRD-1 could no longer locate the females, while males expressing *C. remanei* SRD-*1 or chimeric SRD-1* could still find mates within relatively short periods (∼3 ± 0.9 and ∼6 ± 2.1 minutes for 15 mating pairs; ∼3 ± 1.02 and ∼6 ± 3 minutes for 10 mating pairs; ∼4 ± 2.3 and ∼11 ± 3.8 minutes for 5 mating pairs). However, in experiments with only one male and one female, none of the *C. elegans* transgenic males were able to engage in mating, regardless of the receptor variants expressed. This suggests that SRD-1 plays a pivotal role in facilitating mating system-specific sexual traits in *Caenorhabditis* males, likely due to the polymorphisms within its cytoplasmic domain. Alternative receptor variants should have been eliminated over time, had they not aided in the stabilization of androdioecy. Subsequent investigations into the functional relevance of these polymorphic changes led us to confirm that the cytoplasmic domain variants alter the downstream interacting partner of the receptor (Kannan and Chow, unpublished). Characterizing the downstream interacting partners of the distinct polymorphic cytoplasmic domain of SRD-1 in each species may elucidate the varying cellular events associated with the distinct behavioral modifications.

**Figure 3:**
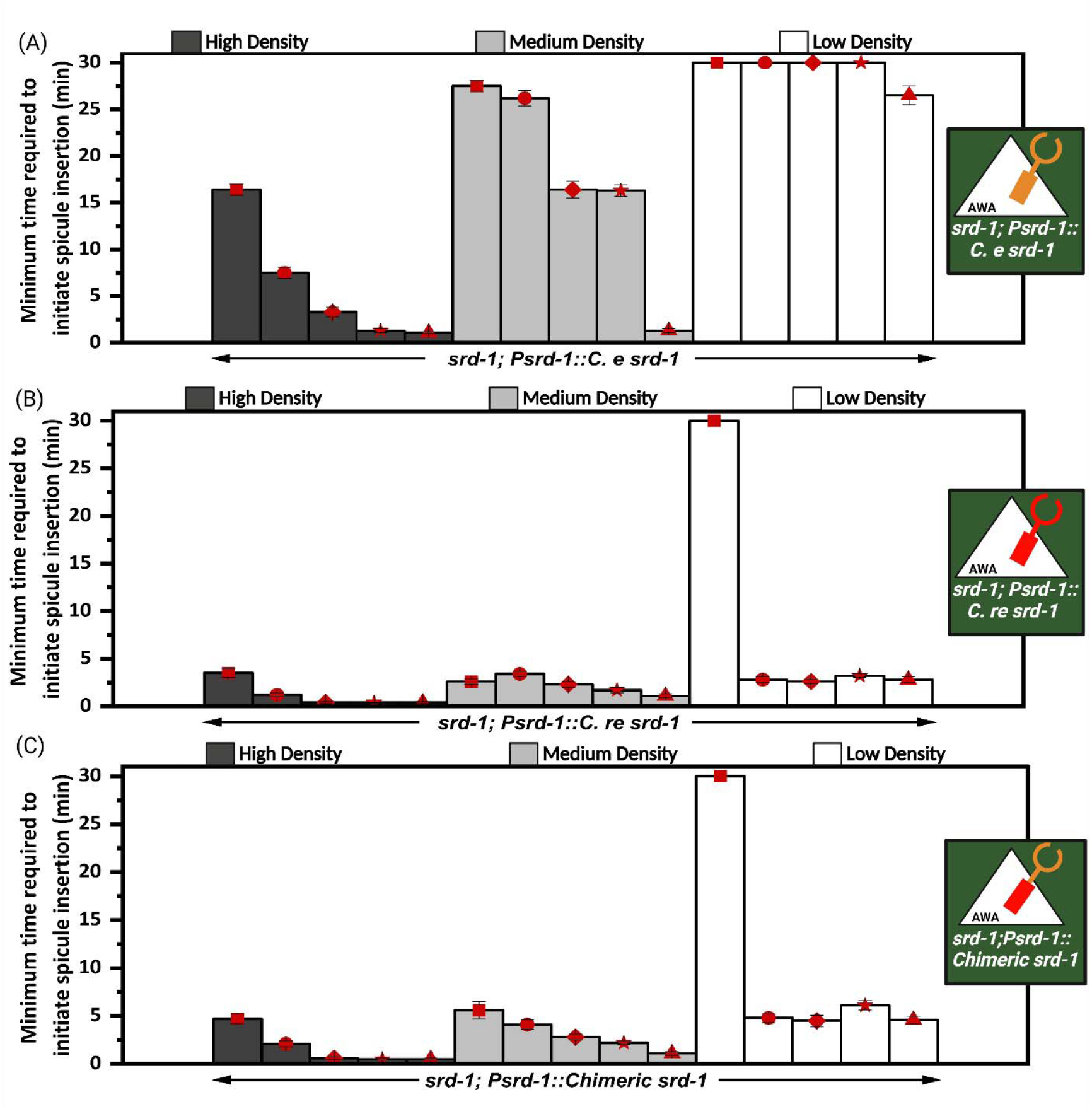
Minimum time for *C. elegans srd-1* mutant males expressing (A) *C. elegans* SRD-1, (B) *C. remanei* SRD-1, and (C) Chimeric SRD-1 to initiate vulval prodding on a *C. remanei* female was recorded under varying density conditions (square: 1 pair; circle: 5 pairs; diamond: 10 pairs; star: 15 pairs; triangle: 20 pairs; *n* = 15).

### Critical facilitating factors for dioecy-androdioecy transition

Ecological conditions favoring the establishment and maintenance of androdioecy are infrequent in natural environments.^30,31^ We propose that contrary to the prevalent belief of ecological factors such as low population density favoring hermaphroditism;^11^ evolution of hermaphroditism could have occurred even in community of higher population density if males and females did not frequently encounter each other. Ancestral female variants exhibiting reduced sexual allure, together with ancestral males that misperceive the availability of mating partners, lead to dramatic reduction of mating opportunities. As a result, a reproductive equilibrium of an emerging androdioecious population dominated by hermaphrodites can be assured. The emergence of hermaphroditism likely had cryptic female variants exhibiting aberrant spermatogenesis and/or oogenesis; mutations compromising self-sperm production and activation are insufficient to ensure a 100% probability of self-fertilization and offspring viability.^32^ Unfit variants were likely purged over generations, a process where elimination of unfit female variants is counterbalanced by abundant progeny in each generation. As such, offsprings with optimal genetic combinations can support a healthy self-fertilizing hermaphroditic population. Despite producing a large number of progenies per generation, this process would have required more than hundreds of generations to fully restore the fitness of their ancestral outbreeding population.^33^ Retention of a small number of males in these populations, due to the residual non-disjunction events, ensures low level of genetic variation created within a confined population thereby mitigating inbreeding depression.^7^ Occasional mating with the residual males instead of complete elimination of males could have played a key role in sustaining a healthy hermaphroditic population as it expands and is stabilized. Our results imply that as androdioecy evolved from dioecious ancestors, androdioecious males continue to express downstream components of SRD-1, which may facilitate dioecious male-like behaviors, i.e., weak habituation and strong female exploration, as an evolutionary vestige.

Based on our and others’ findings, we propose that establishment of hermaphroditism necessitates five key requirements: (1) self-sperm generation, (2) self-sperm activation, (3) diminished sex pheromone potency, (4) strong habituation to sex pheromones, and (5) decreased male mate-seeking behavior. Requirements 1, 2, and 3 are linked to physiological changes in female reproduction, while 4 and 5 pertain to modifications in chemosensory communication centered on males. All the above events very likely occurred in parallel to allow hermaphroditism to gain dominance in the ancestral population. Our data reveal that males from *C. briggsae* and *C. tropicalis* exhibit androdioecious male characteristics resembling *C. elegans* (Sup Fig 1&2). SRD-1 receptors in these species also show significant polymorphisms in their cytoplasmic domain (Sup Fig 3), suggesting that the observed male traits in *C. briggsae* and *C. tropicalis* could also converge on SRD-1. Nonetheless, our study indicates that the evolutionary transition towards self-fertilization in sparse populations^34–36^ may not be obligatorily dependent on a context of low population density. This evolutionary process might also arise from alterations in chemosensory processing, resulting in behavioral adaptations associated with these androdioecious males.

## Author Contributions

KLC conceptualized and supervised the entire project, while HK performed the experiments, conducted data analysis, and wrote the manuscript in consultation with KLC.

## Acknowledgements

We thank former lab members Gus Chan and Olivia Wan for their GC-MS analysis of dioecious and androdioecious sex pheromone profiles and for constructing the SRD-1 clones used in this study. *C. sinica, C. wallecei*, and *C. tropicalis* were kindly provided by Dr. Zhongying Zhao from Hong Kong Baptist University. Other worm strains were sourced from the *Caenorhabditis* Genetics Center. This project was supported by Research Grants Council, Hong Kong CERG 660508 and 660513. We acknowledge the use of OriginPro, Version 2022b (64-bit), SR1.

OriginLab Corporation, Northampton, MA, USA, for data analysis and visualization. Figures were created using BioRender.com.

---

**Supplementary Figure 1:**
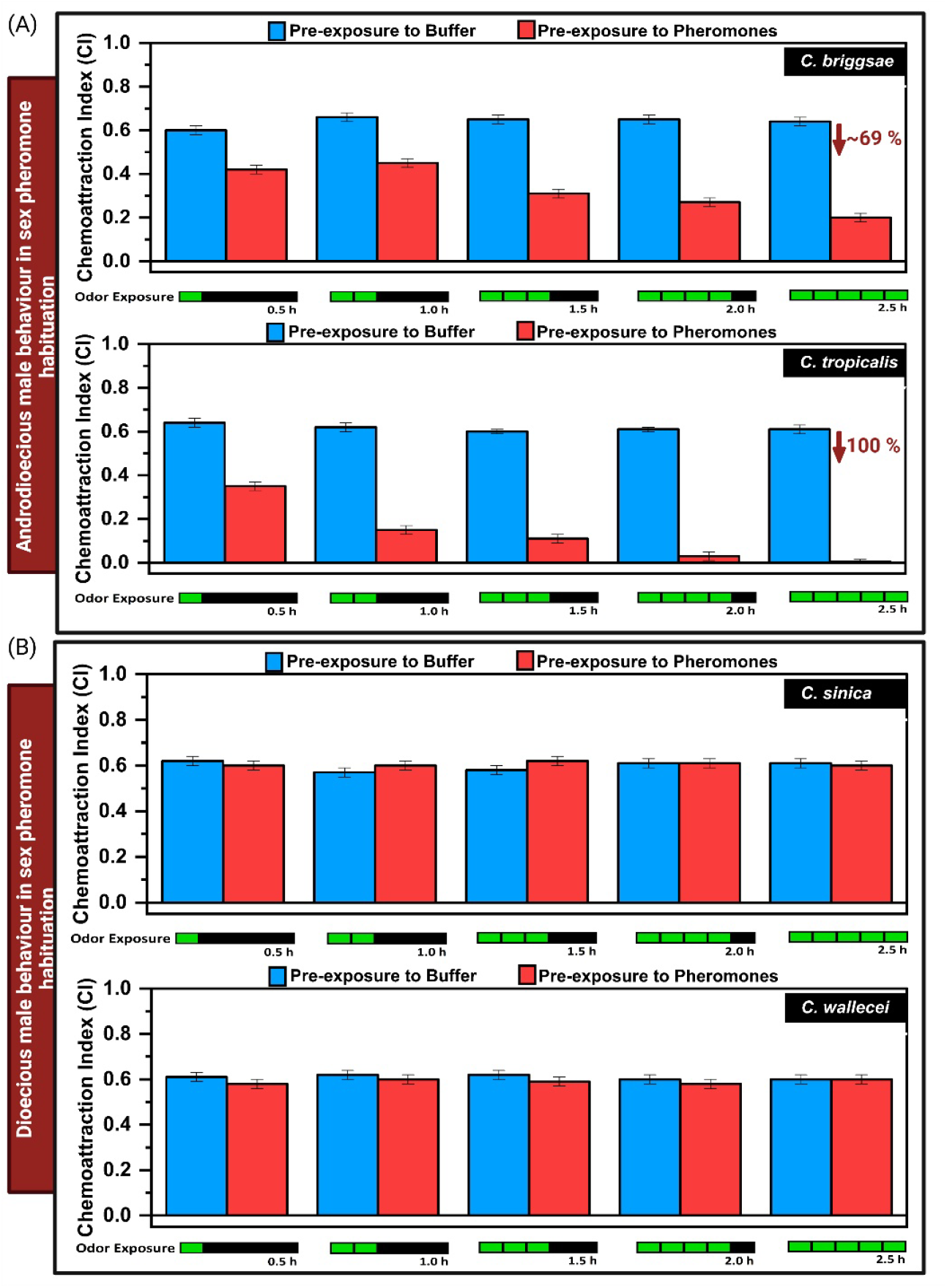
Olfactory habituation was evaluated by measuring responses of androdioecious (A) and dioecious (B) males to *C. remanei* sex pheromones following pre-exposure to pheromones or M9 buffer (*n* = 400 males).

**Supplementary Figure 2:**
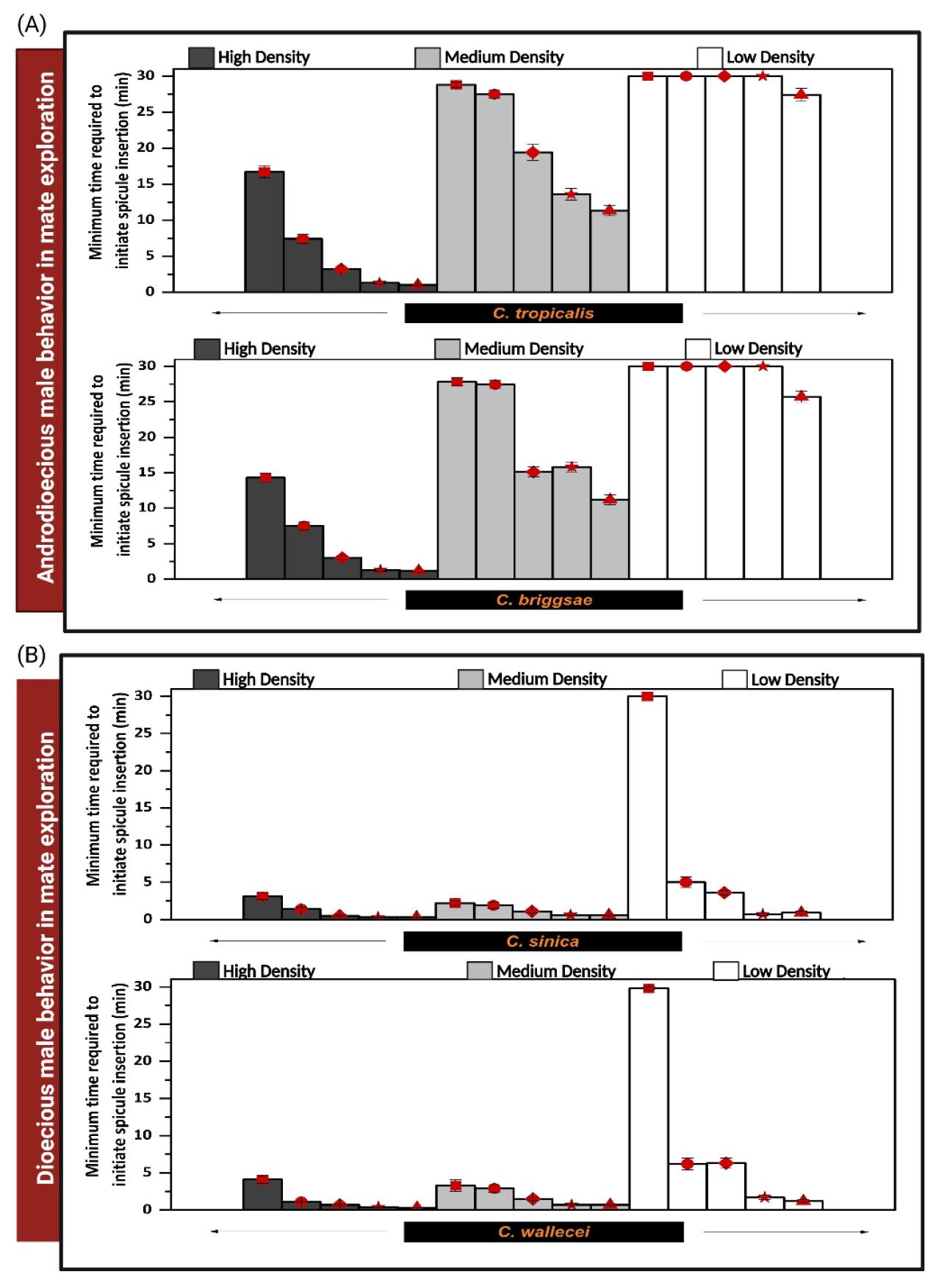
Minimum time for androdioecious (A) and dioecious (B) males to initiate vulval prodding on a *C. remanei* female was recorded under varying density conditions (square: 1 pair; circle: 5 pairs; diamond: 10 pairs; star: 15 pairs; triangle: 20 pairs; *n* = 15).

**Supplementary Figure 3:**
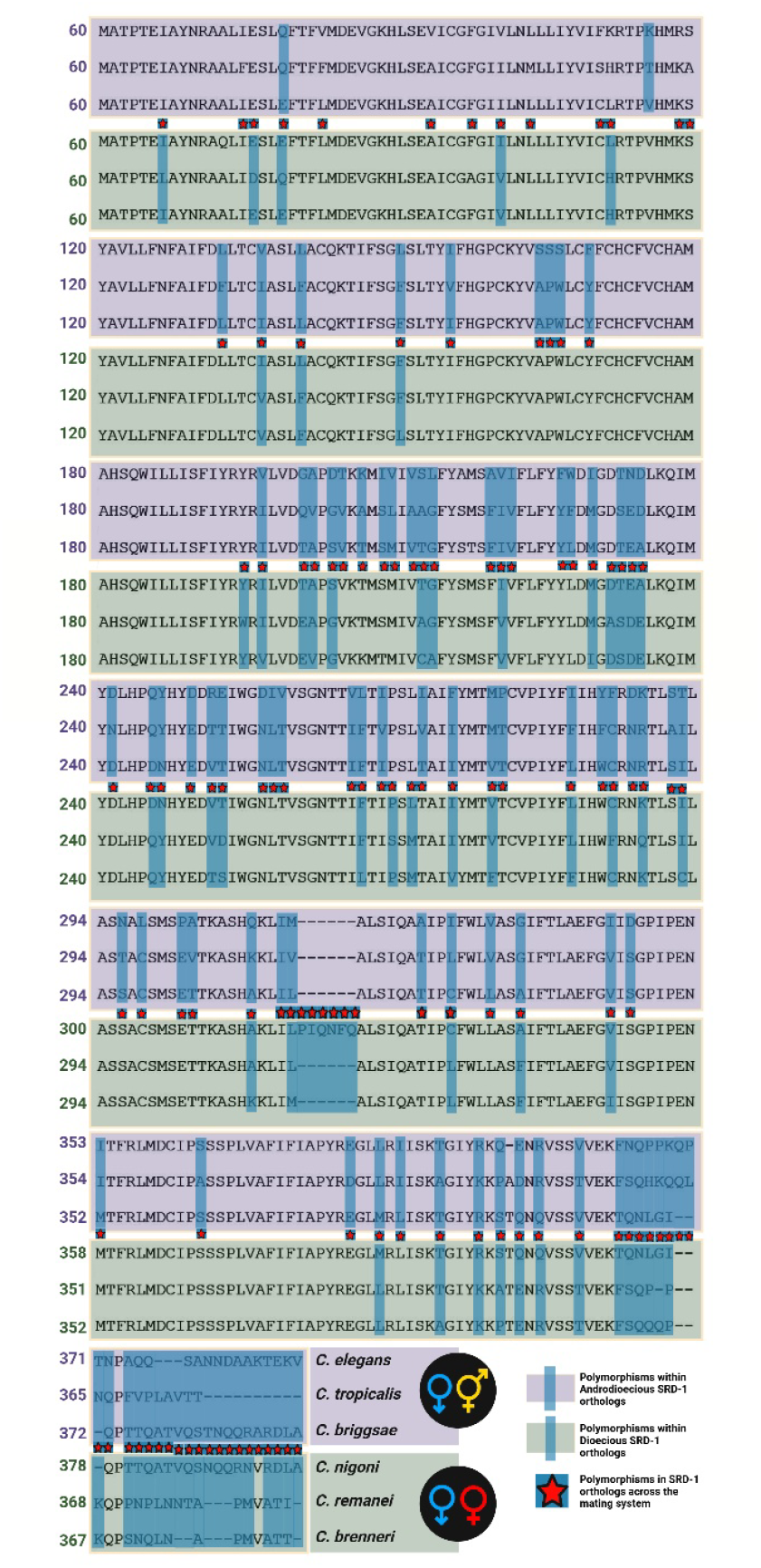
Comparative protein sequence alignment of SRD-1 orthologs was performed using Clustal Omega and visualized with BioRender.

**Supplementary Figure 4:**
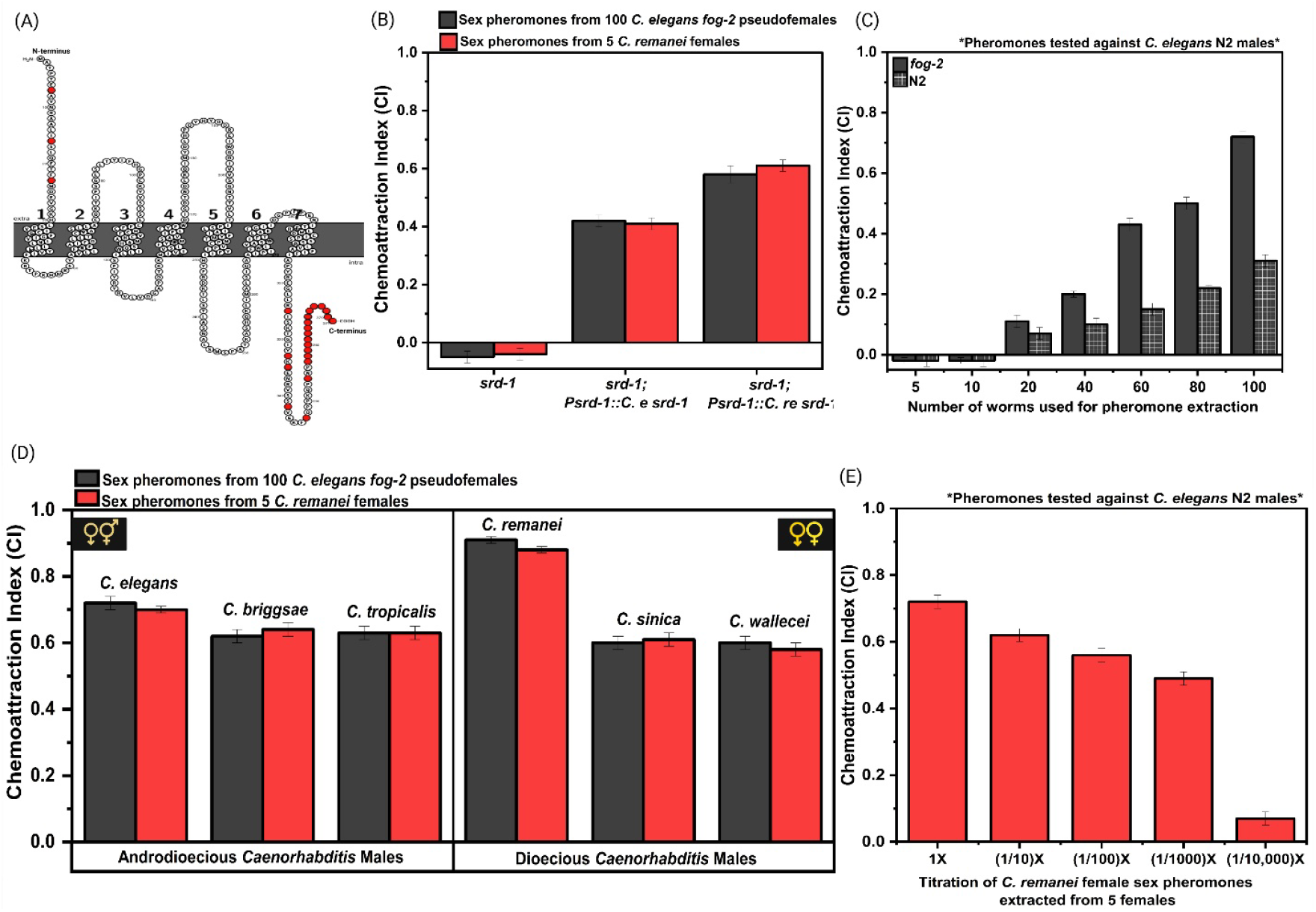
(A) Protein sequence alignment of SRD-1 orthologs was performed using Clustal Omega, with polymorphisms highlighted by red circles and visualized using Protter; (B) *C. elegans srd-1* mutant males expressing SRD-1 orthologs from *C. elegans* and *C. remanei* under the native *srd-1* promoter were tested for their response to sex pheromones extracted from *C. remanei* females and *C. elegans fog-2* pseudo-females (*n* = 400); (C) Sex pheromones were extracted from varying numbers of one-day-old N2 and *fog-2 C. elegans* hermaphrodites and tested against *C. elegans* N2 males (*n* = 400); (D) Males from various dioecious and androdioecious *Caenorhabditis* species were assessed for their behavioral responses to sex pheromones extracted from *C. rem*anei females and *C. elegans fog-2* pseudo-females (*n* = 400); (E) Sex pheromones extracted from five one-day-old virgin *C. remanei* females were titrated and tested against *C. elegans* N2 males (*n* = 400).

